# PulmonDB: a curated lung disease gene expression database

**DOI:** 10.1101/726745

**Authors:** Ana B. Villaseñor-Altamirano, Marco Moretto, Alejandra Zayas-Del Moral, Mariel Maldonado, Adrián Munguía-Reyes, Yair Romero, Jair. S. García-Sotelo, Luis Alberto Aguilar, Aldana-Assad Oscar, Kristof Engelen, Moisés Selman, Julio Collado-Vides, Yalbi I. Balderas-Martínez, Alejandra Medina-Rivera

**Author notes:** To whom correspondence should be addressed to Alejandra Medina-Rivera., Correspondence should also be addressed to Yalbi I. Balderas-Martínez., Correspondence should also be addressed to Julio Collado-Vides.

## Abstract

Chronic Obstructive Pulmonary Disease (COPD) and Idiopathic Pulmonary Fibrosis (IPF) have contrasting clinical and pathological characteristics, and interesting whole-genome transcriptomic profiles. However, data from public repositories are difficult to reprocess and reanalyze. Here we present PulmonDB, a web-based database (http://pulmondb.liigh.unam.mx/) and R library that facilitates exploration of gene expression profiles for these diseases by integrating transcriptomic data and curated annotation from different sources. We demonstrated the value of this resource by presenting the expression of already well-known genes of COPD and IPF across multiple experiments and the results of two differential expression analyses in which we successfully identified differences and similarities. With this first version of PulmonDB, we create a new hypothesis and compare the two diseases from a transcriptomics perspective.

## Background

A common way to study diseases is by using transcriptomic analysis, which can reveal components of the genome that are active and help us understand which biological processes are affected[1]. Over the years, transcriptomic profiles have been compiled and published in public repositories such as Gene Expression Omnibus (GEO)[2,3] and ArrayExpress[4]. Having a way to compare transcriptomic data from Chronic Obstructive Pulmonary Disease (COPD) and Idiopathic Pulmonary Fibrosis (IPF) will help identify common and distinct molecular mechanisms for these two diseases. However, an overwhelming task is to integrate high-throughput data from public repositories, because of platform differences (resulting in batch effects), heterogeneous experimental conditions and the lack of uniformity on experimental annotations. Wang *et al*. reviewed different approaches in which they discussed tools such as GEO2R[5], ScanGEO[6], ImaGEO[7], BioJupies[8]. These tools reuse public data, reanalyze it consistently, and integrate additional data. Even with these available tools, performing meta-analyses is still challenging[9]. In particular, for COPD and IPF, because information from only a few experiments is available in these resources, such an analysis requires manual annotation by the user or inclusion of only curated GEO Datasets (also referred as GDS), and none of them integrates microarray and RNA-Seq data, to our knowledge.

Therefore, we created a curated gene expression lung disease database, PulmonDB, to organize the currently large amount of expression data for both COPD and IPF. To accomplish this task, we used COMMAND>_, a web application previously used to create two successful transcriptomic compendia: one for bacterial genomes, COLOMBOS[10,11], and the second for grapevine VESPUCCI[12]. While there are other chronic respiratory diseases, such as asthma, cystic fibrosis, and pulmonary hypertension association, among others, given the biological similarities between COPD and IPF we decided to focus the first version of PulmonDB on these two diseases. We integrated transcriptomic experiments from different sources and their curated annotations, created an online web resource to facilitate the exploration of gene expression profiles for COPD and IPF creating a new hypotheses, and to allow for the identification of co-expression patterns in further analyses.

## Construction and content

### Data sources

In GEO, we searched for gene expression data sets related to COPD and IPF, we used raw data and metadata available. In specific cases, the platform information (.cdf file) was obtained from the Affymetrix website. Additional information, (*e.g.*, clinical data, source of the biological sample), was obtained either from metadata or manually curated from the original papers. We only considered microarray experiments with available raw data and platform information and we only kept a single copy of each sample.For RNA-seq data we searched in Recount2 [13] to obtain raw counts per gene.

PulmonDB is a relational database, implemented in MySQL with lung disease transcriptome measurements, re-annotated platform probes, and manually curated data with a controlled vocabulary designed for lung diseases. Tables were created to describe each feature and to connect the information across experiments, samples, measurements, platforms, genes, and annotated information. Raw and normalized data, microarray probes and mapped regions can be accessed using MySQL queries and an R package (https://github.com/AnaBVA/pulmondb) that allows users to access and download the data in an R environment. The full database scheme is provided in Supplementary Figure 1.

### Compendium creation

The compendium creation process was done as previously described in COLOMBOS and VESPUCCI[10,12]. In brief, after we filtered the public experiments from GEO that were related to COPD or IPF, we selected their GSE (the experiment ID from GEO), and used COMMAND>_[14]. This web application provides a framework to perform the following steps: 1) download data from selected experiments, 2) parse files and store data in database form, 3) probe-to-gene (re)mapping process, 4) sample curation and annotation, 5) selection of references and sample experiments to determine contrasts, 6) homogenization (and normalization) of data, and 7) perform data quality control (Figure 1).

**Figure 1.**
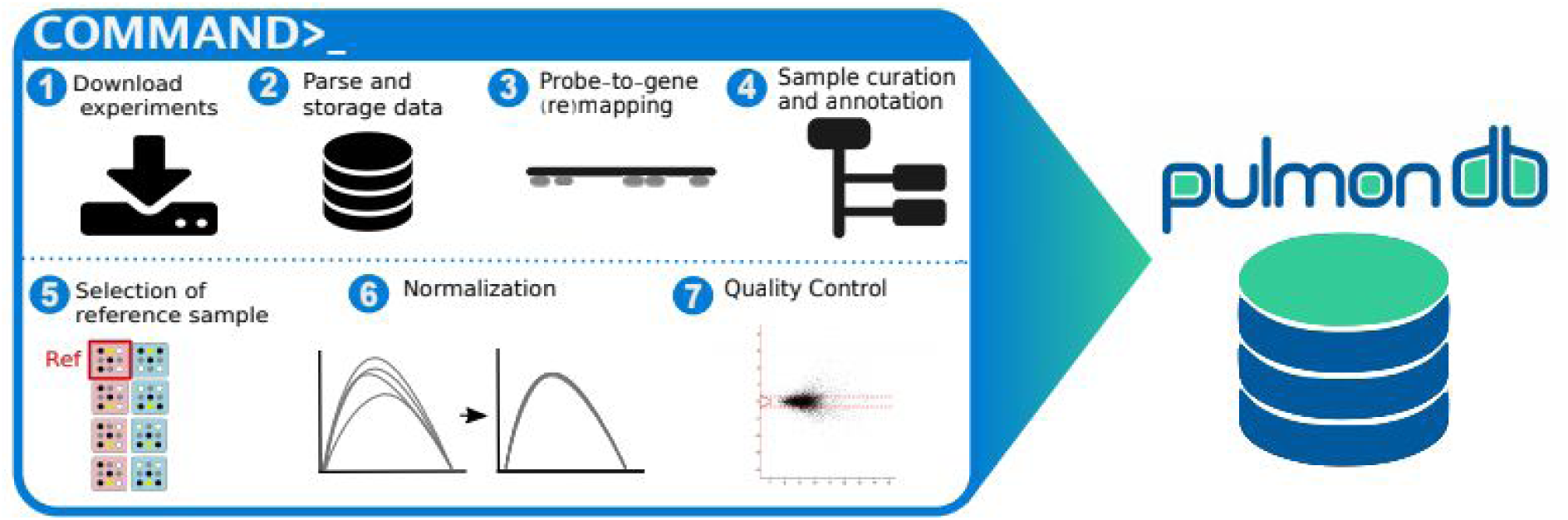
Flow chart of PulmonDB. PulmonDB was created using COMMAND by downloading, parsing and storing COPD and IPF public transcriptomic data into a MySQL database. Then, we remapped microarray probes to establish a uniform gene annotation, and we also created a controlled vocabulary for clinical and biological annotations for each sample. We created contrasts based on the original hypothesis, selecting a sample as the reference. Finally, the data were homogenized and subjected to a quality check.

In more detail, each experiment with a GSE ID, also referred to as a data set, was normalized independently without performing background correction, as explained in [11]. We defined a contrast for each sample with a GSM ID (sample ID from GEO) by using a unique control reference sample per data set. The sample contrast per gene was defined as the log ratio between the expression value in the test condition (*i.e*., IPF, COPD) and the expression value in the reference condition (*i.e*., healthy, untreated, smokers without COPD) (Figure 1, step 5). This gives every comparison an interpretable biological meaning when combined with extensive manual curated annotation. The condition properties describing the contrasts were then structured in a condition-controlled vocabulary tree. Finally, all contrasts were homogenized, resulting in direct comparable log ratios across all experiments; this information then became part of the final compendium of expression data (Supplementary Figure 2).

### PulmonDB uses a controlled vocabulary to describe sample metadata

A controlled vocabulary is required to create databases with homogeneous and standard information. For PulmonDB, we created a controlled vocabulary organized in a hierarchical structure that contains terms to annotate transcriptome experiments in lung diseases. We defined classes describing the main categories and terms that can be found in experiments, with some of them as mandatory features (*e.i.* sample type, sample status, and platform). Some non-IPF or non-COPD diseases were included in the controlled vocabulary because the original experiments used them.

Once the controlled vocabulary was established, each article related to the experiment was manually curated, and whenever it was necessary, new terms were added, making the vocabulary flexible and allowing for the inclusion of other diseases to our database in the future. Complete definitions of the terms are provided in Supplementary Table 1.

### Experiment annotation

Each sample was manually annotated using the controlled vocabulary; when necessary the vocabulary was updated to include new features. The information was curated by experts who reviewed the associated articles and protocols to retrieve data such as age, sex, ancestry, stage of disease or treatment, DLCO (the diffusing capacity of the lung for carbon monoxide, a common functional test), etc., from either GEO or the associated paper.

### Homogenization and quality control

As described before, data homogenization was done with COMMAND>_[11,12]. This step was performed on raw data without background correction, as it has been shown to retrieve more errors[15]. A nonlinear model was applied to homogenize raw data. We used RMA Quantile for Affymetrix samples, and loess fit for the other platforms. The next step was to summarize probes per transcript using RMA median polish summary from Affymetrix or with data averaged across replicates for the other platforms. After performing the homogenization step, low-quality microarrays were identified using MA plots and histograms of raw and homogenized data.

### Website implementation

PulmonDB has a web interface that uses Clustergrammer (https://clustergrammer.readthedocs.io/index.html)[16] to visualize gene expression contrasts. Clustergrammer has a frontend in javascript and a backend in python supporting an interactive web application for gene expression exploration. The PulmonDB web interface requires one or several GSE identifiers and more than two gene names to generate interactive heatmaps.

In addition, Clustergrammer is connected with EnrichR (http://amp.pharm.mssm.edu/Enrichr/)[17], an integrative web application tool for enrichment analysis that helps the user explore not only potentially differentiated genes but also enriched pathways, facilitating the discovery of transcriptomic signature patterns in lung diseases or related phenotypes.

### COPD and IPF comparative analysis

We used limma 3.40.0 in an Rstudio environment 3.6.0 for our comparative analyses, and the GSE ID was included in the linear model. Then, two contrasts were created: 1) “COPD – IPF”, for obtaining differentially expressed genes between COPD and IPF, and 2) “(COPD + IPF)/2 – CONTROL”, for genes similarly expressed between COPD/IPF and CONTROL. Differential gene expression analyses were adjusted for multiple testing using false discovery rate (FDR) method, also referred to Benjamini & Hochberg adjustment. We applied a cutoff of adjusted *p*-value of < 0.05, and after sorting based on the log fold change, the top 20 genes were obtained.

## Utility and discussion

### PulmonDB a curated gene expression lung disease database

Our database has a total of 75 GSEs, corresponding to 4931 unique preprocessed GSMs that used 27 different platforms or GPLs (platform ID from GEO) (Figure 2C). PulmonDB contains different sample types, because we searched for human gene expression experiments related to COPD and IPF without any restriction. Lung biopsies account for 25.3% of samples and 43.7% are blood samples. However, different cell types can be found in PulmonDB: some of them are primary cells (*e.i.* alveolar macrophages, fibroblasts, alveolar epithelial cells, etc.) and others are cell lines (*e.i.* A549) (Figure 2A). Of the samples, 42.3% correspond to COPD, 33.4% to healthy/control, 13.6% to IPF, 6.2% to match tissue/control, and 4.4% to other diseases (Figure 2B and Supplementary Table 2). PulmonDB is a curated gene expression database of human lung diseases, with RNA-seq and microarray data from different platforms that have been uniformly preprocessed and manually curated to add sample and experiment information. In addition, we developed a website to access and visualize homogenized data (http://pulmondb.liigh.unam.mx/), and we also developed an R package (https://github.com/AnaBVA/pulmondb to download curated annotation and preprocessed data that can be used for further analysis in the R environment.

**Figure 2.**
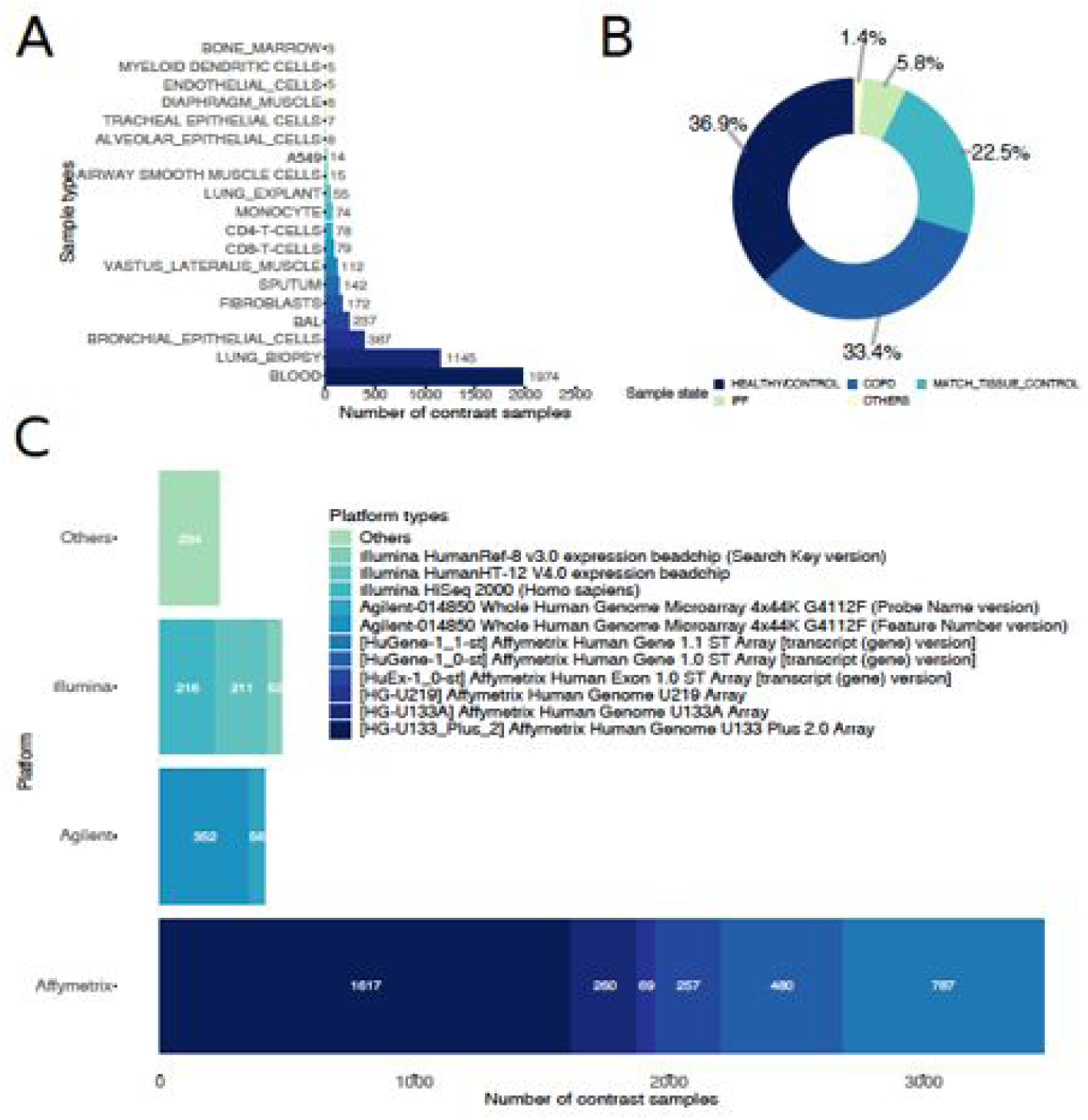
Summary of PulmonDB. A) Number of contrast samples in PulmonDB per biological sample type. B) Proportions of sample states found in PulmonDB. The color key below the pie chart shows the sectors for COPD patients, healthy/controls, IPF patients, match_tissue_controls (nontumor cancer patients), and other diseases (such as asthma). C) Number of contrast samples measured using each platform (clustered by using Affymetrix, Agilent, Illumina and other platforms with fewer samples).

Although, there are other resources that reuse, and reanalyze GEO data using web interfaces[9], those tools are not specialized for lung diseases. Their limitations include the need for previous manual curation in each analysis, and they only consider a small number of COPD and IPF experiments due to the fact that only curated GEO data are used. We designed a web interface that enables data exploration and visualization to facilitate lung disease analysis. This interface uses Clustergrammer[16] to visualize gene expression values and the creation of interactive heatmaps that allow data exploration. In addition, Clustergrammer is connected to EnrichR[17], which allows pathway enrichment analysis. All these features together should help to generate new hypotheses about the pathologies of lung diseases to perform exploratory analyses, to visualize specific gene expression across public experiments for comparing results, and to generate new insights based on different data sets.

### PulmonDB can recapitulate gene expression patterns expected in COPD and IPF

To show that PulmonDB can be used to recapitulate previously reported knowledge regarding COPD and IPF biology, we performed a literature search and manually selected relevant genes for each disease. We selected 19 genes related to IPF (not necessarily associated with gene expression in lung tissues) to visualize their gene expression: CCL18[18], CXCL12[19], CXCL13[20], collagens (COL1A1, COL1A2, COL3A1, COL5A2, COL14A1)[21], DSP[22], FAS[23], IL-8[24], MMP1[25], MMP2[26], MMP7[25], MUC5B[22], SPP1[27], PTGS2[28], TGFB1[29] and THY1[30]. Then, we selected eight IPF experiments performed with lung tissue biopsy samples (GSE32537, GSE21369, GSE24206, GSE94060, GSE72073, GSE35145, GSE31934), and using the PulmonDB website, we created a heatmap with the gene expression patterns and observed that the hierarchical clustering of these data separates IPF and control data sets (Figure 3A, green and grey clusters at the bottom). For COPD, we curated 16 genes from the literature that were deemed relevant to this disease: HHIP[31,32], CFTR[33,34], PPARG[35], SERPINA1[36,37], JUN[38], FAM13A[39], MYH10[38], CHRNA5[40], JUND[38], JUNB[38], TNF[37], MMP9[37], MMP12[37], CHRNA3[40], TGFBR3[35], and GATA2[35]. We selected five experiments (GSE27597, GSE37768, GSE57148, GSE8581, GSE1122) performed on lung tissue biopsy samples from COPD patients and controls. Our hierarchical clustering analysis of the expression profiles using the PulmonDB interface allowed us to cluster patients and controls into two different groups (Figure 3B), similar to the case of IPF. In conclusion, PulmonDB not only helps to recapitulate previously published work (Supplementary Figure 3) but also helps to verify their expression stability across experiments. This may help to analyze concordance in different experiments, contrast study results, show implications of using different control groups, etc. We believe this resource can be used to drive, make decisions and support new hypotheses in experimental laboratories for studying molecular or cellular disease mechanisms.

**Figure 3.**
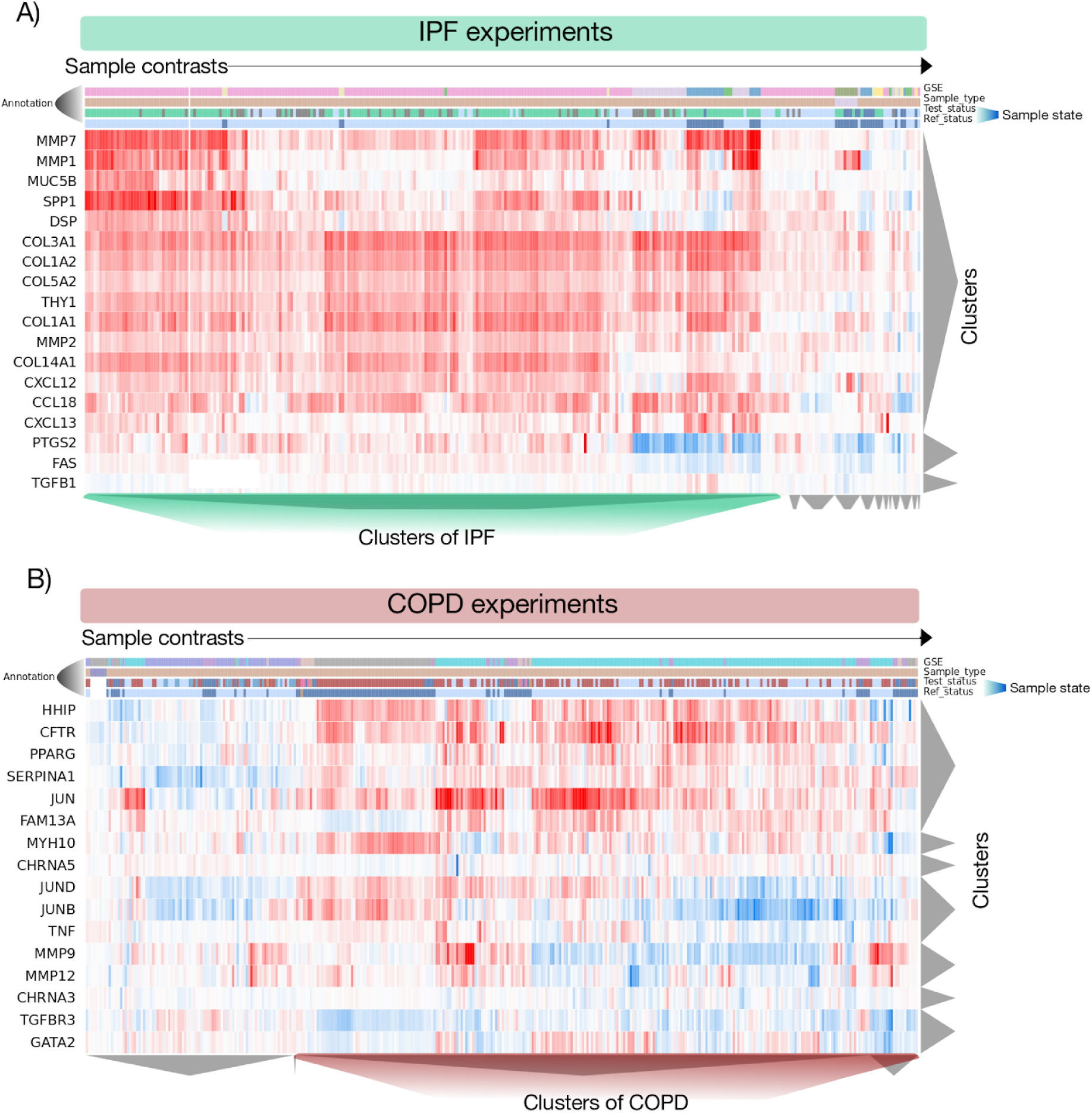
IPF and COPD well-known disease-associated genes. In both heatmaps, rows are genes and columns are sample contrasts. Both were hierarchically clustered. The first annotation row represents their GSE IDs. The second annotation row is the sample type, LUNG_BIOPSY samples, in light brown. The third and the fourth annotation rows are sample states, the third annotation row represents the test state and the fourth annotation row is the reference state. A) IPF genes reported being relevant in the literature (CCL18[18], CXCL12[19], CXCL13[20], COL1A1, COL1A2, COL3A1, COL5A2, COL14A1[21], DSP[22], FAS[23], IL-8[24], MMP1[25], MMP2[26], MMP7[25], MUC5B[22], SPP1[27], PTGS2[28], TGFB1[29] and THY1[30]). The IPF experiments selected were GSE32537 (pink), GSE21369 (purple), GSE24206 (blue), GSE94060 (grass-green), GSE72073 (lemon yellow), GSE35145 (green), and GSE31934 (yellow). The third and the fourth annotation rows are sample states: light blue, MATCH_TISSUE_CONTROL; dark blue, HEALTHY/CONTROL; turquoise, IPF samples; and grey, NON_IPF_ILD. B) COPD genes reported being relevant in the literature (HHIP[31,32], CFTR[33,34], PPARG[35], SERPINA1[36,37], JUN[38], FAM13A[39], MYH10[38], CHRNA5[40], JUND[38], JUNB[38], TNF[37], MMP9[37], MMP12[37], CHRNA3[40], TGFBR3[35], and GATA2[35]). The COPD experiments selected were GSE27597, GSE37768, GSE57148, GSE8581, and GSE1122. The third and the fourth annotation rows are sample states: light blue, MATCH_TISSUE_CONTROL; dark blue, HEALTHY/CONTROL; red, COPD samples.

### Differences and similarities in COPD and IPF

PulmonDB can be used not only to replicate previous knowledge but also to provide a framework to test new hypotheses. In this context, we set out to investigate the differences and similarities between COPD and IPF in lung tissue when compared to samples from healthy individuals (Figure 4A). Using PulmonDB in the R environment, we selected contrasts where the sample was annotated as lung biopsy and the reference status as HEALTHY/CONTROLs (GSE52463, GSE63073, GSE1122, GSE72073, GSE24206, GSE27597, GSE29133, GSE31934, GSE37768) (Figure 4B), and then using limma[41] we assessed differential gene expression between COPD and IPF, we identified 1781 differentially expressed genes (Supplementary Figure 4). To have a visual representation of the differences between COPD and IPF, we selected the top 20 differentially expressed genes and visualized their expression using the PulmonDB website tool (Figure 4C). We observed that data sets tend to cluster by test status; Figure 4C shows IPF contrasts on the left (turquoise), control contrasts in the middle (blue), and COPD contrasts on the right (red). Genes are clustered in two groups (left panel, y-axis); the first gene group (I) is overexpressed in IPF while it is barely expressed or underexpressed in COPD contrasts. By comparison, the second gene cluster (group II) is overexpressed in COPD contrasts and underexpressed in IPF. To correlate similarities among samples, the 20 top differentially expressed genes were used (Figure 4C, right panel); samples from the same disease group showed higher correlations, and tended to have a null or negative correlation with the HEALTHY/CONTROL and the opposite disease (Figure 4C). For example, FOSB and CXCL2 have opposite behaviors, as both genes are overexpressed in COPD and underexpressed in IPF. FOSB is part of the family of Fos genes that can dimerize with JUN family proteins to form the transcription factor complex AP-1, which is related to COPD[42]. CXCL2 is a chemokine secreted in inflammation that induces chemotaxis in neutrophils[43,44]; these cells are predominant in COPD and they are key mediators in tissue damage[45]. While neutrophils are also important in IPF, we observed their underexpression in this disease.

**Figure 4.**
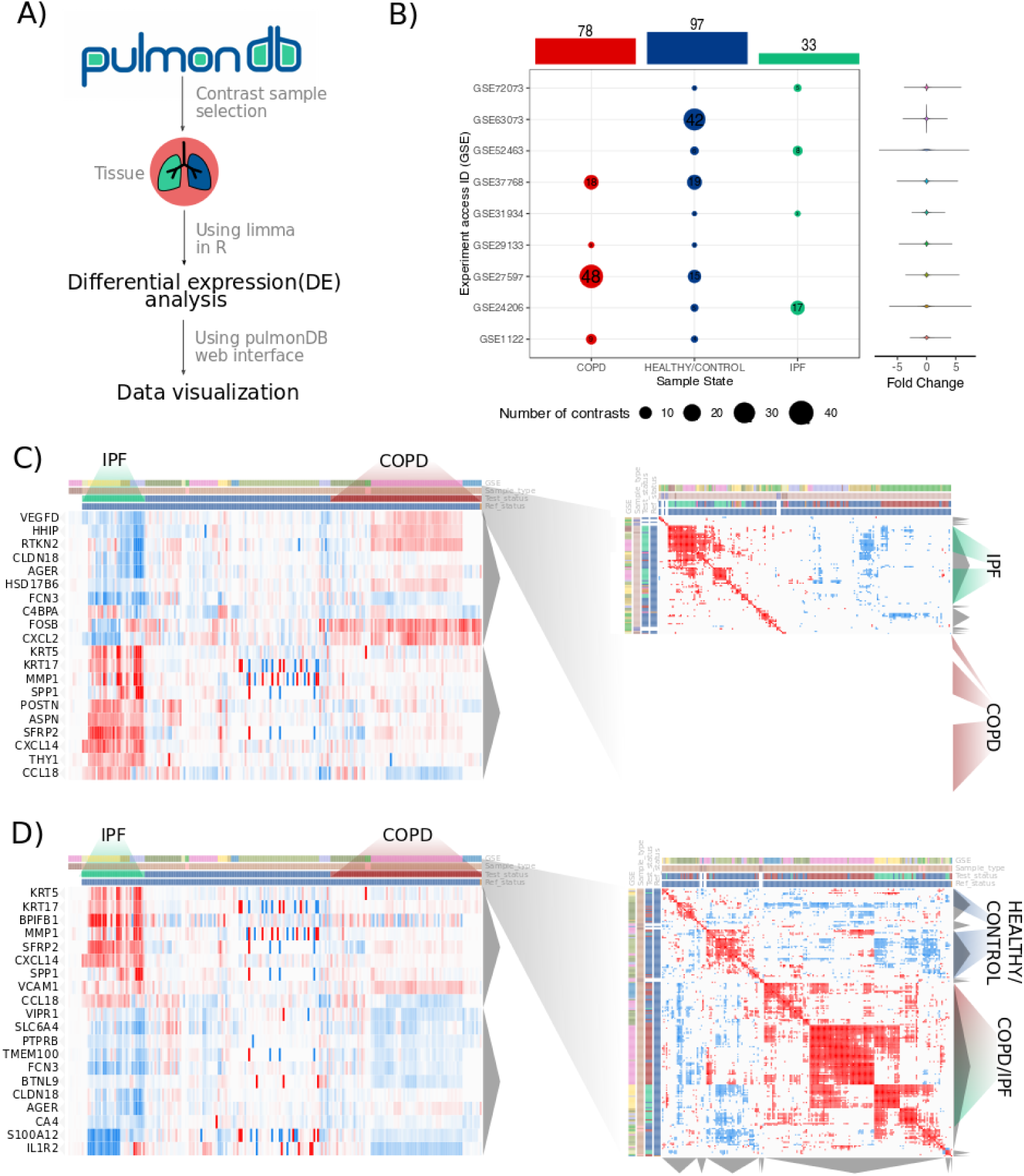
IPF and COPD differentially expressed and similarly expressed genes. A) Flow chart of steps used for COPD and IPF differential expression analysis to evaluate transcriptomic differences and similarities. B) Experiments selected for the analysis, following the criteria of being lung biopsy samples and contrasted with HEALTHY/CONTROL references. The colors represent the sample state: COPD, red; HEALTH/CONTROL, blue; IPF, turquoise. At the top, the bar graph is the total sum of contrasts, rows are the GSE experiments, and each dot is the number of contrasts per experiment from COPD, HEALTHY/CONTROL, or IPF subjects. On the right side, we can see the distributions in volcano plots for all sample contrasts per experiment. C) Differentially expressed genes between COPD and IPF. D) Similar genes between COPD and IPF. In both C) and D), columns are sample contrasts, rows are genes, the first covariate is colored by each corresponding experiment, the second covariate is the sample type (in this case, lung tissue is shown in light brown), the third row is the test status and the fourth is the reference status. Columns are ordered by test status and genes by hierarchical clusterization. The right heatmap is the correlation among sample contrasts, and the covariates are the same.

We also asked the opposite question, *i.e.*, whether we could identify which genes that are shared between these two diseases. We assigned weight to COPD and IPF expression to perform limma contrasts (Figure 4D), which enabled us to identify when a differential expression profile was driven by both diseases. We selected the 20 top differentially expressed genes and visualized their expression patterns using PulmonDB website tool, and we could see that a set of genes was consistently overexpressed or underexpressed in both COPD and IPF. In particular, VCAM1 and FCN3 are differentially expressed in COPD and IPF, with a similar trend in both diseases when compared with HEALTHY/CONTROLs. VCAM1 is the vascular cell adhesion molecule 1 and it is important in the immune response for mediating cellular adhesion in leukocytes[46]; it is overexpressed in these two diseases, suggesting infiltration of immune cells in both pathologies[47,48]. In contrast, FCN3 (or ficolin 3) is underexpressed in both diseases: this gene is a collagen-like protein associated with the innate immune defense, as it activates the lectin complement pathway[49], which has been shown to be important in pulmonary pathologies [50,51].

As a result, PulmonDB assisted our analysis of COPD and IPF analogous and antagonist genes and can thus be used to dissect common molecular mechanisms, because both lung diseases are present under heterogeneous conditions with progressive and irreversible phenotypes mainly caused by smoking and by aging, plus both diseases entail cellular matrix remodeling. Furthermore, the differential gene signatures between COPD and IPF might explain the particularities of each disease.

## Conclusions

PulmonDB can help the scientific community to study which genes have a distinct expression profile in COPD and IPF, explore experiments across technologies and platforms, identify interesting expression patterns across different diseases, generate new hypotheses, and find relationships among clinical or experimental variables. This database also enables comparisons of an updated collection of expression profiles already homogenized for their analyses of specific diseases. Additionally, having different lung diseases (COPD and IPF) in the same database creates the opportunity to observe their similarities and differences. In the future, we aim for PulmonDB to grow and include more diseases. To our knowledge, there is no other resource for transcriptomic analysis focused on the same lung diseases; for this reason, we believe researchers of different backgrounds can use and benefit from the information contained in PulmonDB, its web interface and its package.

We believe that an integrated comparable collection of homogenized values with controlled vocabulary describing biological and technical characteristics will facilitate further comparative analyses, such as the study of profiles in COPD and IPF, exploration of experiments across technologies and platforms, identification of interesting coexpression patterns across different diseases, the generation of new hypotheses, and determination of relationships among clinical or experimental variables.

## PERSPECTIVES

This project sets the foundation to integrate transcriptomics data of other respiratory diseases or related phenotypes and thus facilitates the identification of common and divergent pathways that lead to a pathological state. The PulmonDB platform will be expanded to include other lung diseases.

## AVAILABILITY

PulmonDB is accessible online (http://pulmondb.liigh.unam.mx/) and through an R package (https://github.com/AnaBVA/pulmondb).

## ACKNOWLEDGEMENTS

We are thankful to the colleagues who help us installing and maintaining the PulmonDB server, particularly Luis Alberto Aguilar Bautista and members of Laboratorio Nacional de Visualización Científica Avanzada, Mexico. We thank Alejandra Castillo and Carina Uribe for technical assistance. We thank Mauricio Guzman and Centro Cultural Cine y Arte, particularly Renata Campuzano and Diego Morales for graphical and design assistance. YIB-M acknowledges her research position within the Cátedras CONACyT program. We also acknowledge Catalina Frank, Jose Antonio Alonso and the Manuscript Writing Training Team (CEMAI for its Spanish acronym) of CONACyT for their help with the structure, reviews and constructive criticism of this research paper.

## REFERENCES

1. Qian X, Ba Y, Zhuang Q, Zhong G. RNA-Seq technology and its application in fish transcriptomics. OMICS. 2014;18:98–110.

2. geo. Home - GEO - NCBI [Internet]. [cited 2019 Jul 21]. Available from: https://www.ncbi.nlm.nih.gov/geo/

3. Clough E, Barrett T. The Gene Expression Omnibus Database. Methods Mol Biol. 2016;1418:93–110.

4. EMBL-EBI. ArrayExpress < EMBL-EBI [Internet]. [cited 2019 Jul 21]. Available from: https://www.ebi.ac.uk/arrayexpress/

5. Barrett T, Wilhite SE, Ledoux P, Evangelista C, Kim IF, Tomashevsky M, et al. NCBI GEO: archive for functional genomics data sets--update. Nucleic Acids Res. 2013;41:D991–5.

6. Koeppen K, Stanton BA, Hampton TH. ScanGEO: parallel mining of high-throughput gene expression data. Bioinformatics. 2017;33:3500–1.

7. Toro-Domínguez D, Martorell-Marugán J, López-Domínguez R, García-Moreno A, González-Rumayor V, Alarcón-Riquelme ME, et al. ImaGEO: integrative gene expression meta-analysis from GEO database. Bioinformatics. 2019;35:880–2.

8. Torre D, Lachmann A, Ma’ayan A. BioJupies: Automated Generation of Interactive Notebooks for RNA-Seq Data Analysis in the Cloud. Cell Syst. 2018;7:556–61.e3.

9. Wang Z, Lachmann A, Ma’ayan A. Mining data and metadata from the gene expression omnibus. Biophys Rev. 2019;11:103–10.

10. Moretto M, Sonego P, Dierckxsens N, Brilli M, Bianco L, Ledezma-Tejeida D, et al. COLOMBOS v3.0: leveraging gene expression compendia for cross-species analyses. Nucleic Acids Res. 2016;44:D620–3.

11. Engelen K, Fu Q, Meysman P, Sánchez-Rodríguez A, De Smet R, Lemmens K, et al. COLOMBOS: access port for cross-platform bacterial expression compendia. PLoS One. 2011;6:e20938.

12. Moretto M, Sonego P, Pilati S, Malacarne G, Costantini L, Grzeskowiak L, et al. VESPUCCI: Exploring Patterns of Gene Expression in Grapevine. Front Plant Sci. 2016;7:633.

13. Collado-Torres L, Nellore A, Kammers K, Ellis SE, Taub MA, Hansen KD, et al. Reproducible RNA-seq analysis using recount2. Nat Biotechnol. 2017;35:319–21.

14. Moretto M, Sonego P, Villaseñor-Altamirano AB, Engelen K. First step toward gene expression data integration: transcriptomic data acquisition with COMMAND>_. BMC Bioinformatics. 2019;20:54.

15. Irizarry RA, Hobbs B, Collin F, Beazer-Barclay YD, Antonellis KJ, Scherf U, et al. Exploration, normalization, and summaries of high density oligonucleotide array probe level data. Biostatistics. Department of Biostatistics, Johns Hopkins University, Baltimore, MD 21205, USA.; 2003;4:249–64.

16. Fernandez NF, Gundersen GW, Rahman A, Grimes ML, Rikova K, Hornbeck P, et al. Clustergrammer, a web-based heatmap visualization and analysis tool for high-dimensional biological data. Sci Data. 2017;4:170151.

17. Chen EY, Tan CM, Kou Y, Duan Q, Wang Z, Meirelles GV, et al. Enrichr: interactive and collaborative HTML5 gene list enrichment analysis tool. BMC Bioinformatics. 2013;14:128.

18. Cai M, Bonella F, He X, Sixt SU, Sarria R, Guzman J, et al. CCL18 in serum, BAL fluid and alveolar macrophage culture supernatant in interstitial lung diseases. Respir Med. 2013;107:1444–52.

19. Antoniou KM, Soufla G, Lymbouridou R, Economidou F, Lasithiotaki I, Manousakis M, et al. Expression analysis of angiogenic growth factors and biological axis CXCL12/CXCR4 axis in idiopathic pulmonary fibrosis. Connect Tissue Res. 2010;51:71–80.

20. Vuga LJ, Tedrow JR, Pandit KV, Tan J, Kass DJ, Xue J, et al. C-X-C motif chemokine 13 (CXCL13) is a prognostic biomarker of idiopathic pulmonary fibrosis. Am J Respir Crit Care Med. 2014;189:966–74.

21. Jenkins RG, Simpson JK, Saini G, Bentley JH, Russell A-M, Braybrooke R, et al. Longitudinal change in collagen degradation biomarkers in idiopathic pulmonary fibrosis: an analysis from the prospective, multicentre PROFILE study. Lancet Respir Med. 2015;3:462–72.

22. Allen RJ, Porte J, Braybrooke R, Flores C, Fingerlin TE, Oldham JM, et al. Genetic variants associated with susceptibility to idiopathic pulmonary fibrosis in people of European ancestry: a genome-wide association study. Lancet Respir Med. 2017;5:869–80.

23. Huang SK, Scruggs AM, Donaghy J, Horowitz JC, Zaslona Z, Przybranowski S, et al. Histone modifications are responsible for decreased Fas expression and apoptosis resistance in fibrotic lung fibroblasts. Cell Death Dis. 2013;4:e621.

24. Yang L, Herrera J, Gilbertsen A, Xia H, Smith K, Benyumov A, et al. IL-8 mediates idiopathic pulmonary fibrosis mesenchymal progenitor cell fibrogenicity. Am J Physiol Lung Cell Mol Physiol. 2018;314:L127–36.

25. Rosas IO, Richards TJ, Konishi K, Zhang Y, Gibson K, Lokshin AE, et al. MMP1 and MMP7 as potential peripheral blood biomarkers in idiopathic pulmonary fibrosis. PLoS Med. 2008;5:e93.

26. García-Alvarez J, Ramirez R, Sampieri CL, Nuttall RK, Edwards DR, Selman M, et al. Membrane type-matrix metalloproteinases in idiopathic pulmonary fibrosis. Sarcoidosis Vasc Diffuse Lung Dis. 2006;23:13–21.

27. Pardo A, Gibson K, Cisneros J, Richards TJ, Yang Y, Becerril C, et al. Up-regulation and profibrotic role of osteopontin in human idiopathic pulmonary fibrosis. PLoS Med. 2005;2:e251.

28. Parra ER, Lin F, Martins V, Rangel MP, Capelozzi VL. Immunohistochemical and morphometric evaluation of COX 1 and COX-2 in the remodeled lung in idiopathic pulmonary fibrosis and systemic sclerosis. J Bras Pneumol. 2013;39:692–700.

29. Martinez FJ, Collard HR, Pardo A, Raghu G, Richeldi L, Selman M, et al. Idiopathic pulmonary fibrosis. Nat Rev Dis Primers. 2017;3:17074.

30. Sanders YY, Pardo A, Selman M, Nuovo GJ, Tollefsbol TO, Siegal GP, et al. Thy-1 promoter hypermethylation: a novel epigenetic pathogenic mechanism in pulmonary fibrosis. Am J Respir Cell Mol Biol. 2008;39:610–8.

31. Zhou X, Baron RM, Hardin M, Cho MH, Zielinski J, Hawrylkiewicz I, et al. Identification of a chronic obstructive pulmonary disease genetic determinant that regulates HHIP. Hum Mol Genet. 2012;21:1325–35.

32. Chang W-A, Tsai M-J, Jian S-F, Sheu C-C, Kuo P-L. Systematic analysis of transcriptomic profiles of COPD airway epithelium using next-generation sequencing and bioinformatics. Int J Chron Obstruct Pulmon Dis. 2018;13:2387–98.

33. Rab A, Rowe SM, Raju SV, Bebok Z, Matalon S, Collawn JF. Cigarette smoke and CFTR: implications in the pathogenesis of COPD. Am J Physiol Lung Cell Mol Physiol. 2013;305:L530–41.

34. Campbell JD, McDonough JE, Zeskind JE, Hackett TL, Pechkovsky DV, Brandsma C-A, et al. A gene expression signature of emphysema-related lung destruction and its reversal by the tripeptide GHK. Genome Med. 2012;4:67.

35. Hedström U, Hallgren O, Öberg L, DeMicco A, Vaarala O, Westergren-Thorsson G, et al. Bronchial extracellular matrix from COPD patients induces altered gene expression in repopulated primary human bronchial epithelial cells. Sci Rep. 2018;8:3502.

36. Lackey L, McArthur E, Laederach A. Increased Transcript Complexity in Genes Associated with Chronic Obstructive Pulmonary Disease. PLoS One. 2015;10:e0140885.

37. Kotnala S, Tyagi A, Muyal JP. rHuKGF ameliorates protease/anti-protease imbalance in emphysematous mice. Pulm Pharmacol Ther. 2017;45:124–35.

38. Kim WJ, Lim JH, Lee JS, Lee S-D, Kim JH, Oh Y-M. Comprehensive Analysis of Transcriptome Sequencing Data in the Lung Tissues of COPD Subjects. Int J Genomics Proteomics. 2015;2015:206937.

39. Yun JH, Morrow J, Owen CA, Qiu W, Glass K, Lao T, et al. Transcriptomic Analysis of Lung Tissue from Cigarette Smoke-Induced Emphysema Murine Models and Human Chronic Obstructive Pulmonary Disease Show Shared and Distinct Pathways. Am J Respir Cell Mol Biol. 2017;57:47–58.

40. Matsson H, Söderhäll C, Einarsdottir E, Lamontagne M, Gudmundsson S, Backman H, et al. Targeted high-throughput sequencing of candidate genes for chronic obstructive pulmonary disease. BMC Pulm Med. 2016;16:146.

41. Ritchie ME, Phipson B, Wu D, Hu Y, Law CW, Shi W, et al. limma powers differential expression analyses for RNA-sequencing and microarray studies. Nucleic Acids Res. 2015;43:e47.

42. Mroz RM, Holownia A, Chyczewska E, Braszko JJ. Chronic obstructive pulmonary disease: an update on nuclear signaling related to inflammation and anti-inflammatory treatment. J Physiol Pharmacol. 2008;59 Suppl 6:35–42.

43. Kim D, Haynes CL. Neutrophil chemotaxis within a competing gradient of chemoattractants. Anal Chem. 2012;84:6070–8.

44. Larsson K. Aspects on pathophysiological mechanisms in COPD. J Intern Med. 2007;262:311–40.

45. Hoenderdos K, Condliffe A. The neutrophil in chronic obstructive pulmonary disease. Am J Respir Cell Mol Biol. 2013;48:531–9.

46. Ley K, Huo Y. VCAM-1 is critical in atherosclerosis. J Clin Invest. 2001;107:1209–10.

47. Nakao A, Hasegawa Y, Tsuchiya Y, Shimokata K. Expression of cell adhesion molecules in the lungs of patients with idiopathic pulmonary fibrosis. Chest. 1995;108:233–9.

48. Davis BB, Shen Y-H, Tancredi DJ, Flores V, Davis RP, Pinkerton KE. Leukocytes are recruited through the bronchial circulation to the lung in a spontaneously hypertensive rat model of COPD. PLoS One. 2012;7:e33304.

49. Garred P, Honoré C, Ma YJ, Munthe-Fog L, Hummelshøj T. MBL2, FCN1, FCN2 and FCN3-The genes behind the initiation of the lectin pathway of complement. Mol Immunol. 2009;46:2737–44.

50. Pandya PH, Wilkes DS. Complement system in lung disease. Am J Respir Cell Mol Biol. 2014;51:467–73.

51. Eisen DP. Mannose-binding lectin deficiency and respiratory tract infection. J Innate Immun. 2010;2:114–22.

52. Fujino N, Ota C, Takahashi T, Suzuki T, Suzuki S, Yamada M, et al. Gene expression profiles of alveolar type II cells of chronic obstructive pulmonary disease: a case-control study. BMJ Open [Internet]. 2012;2. Available from: http://dx.doi.org/10.1136/bmjopen-2012-001553

53. Golpon HA, Coldren CD, Zamora MR, Cosgrove GP, Moore MD, Tuder RM, et al. Emphysema lung tissue gene expression profiling. Am J Respir Cell Mol Biol. 2004;31:595–600.

