## SupplementaryTablesFigures for "PulmonDB: a curated lung disease gene expression database"

### SUPPLEMENTARY DATA

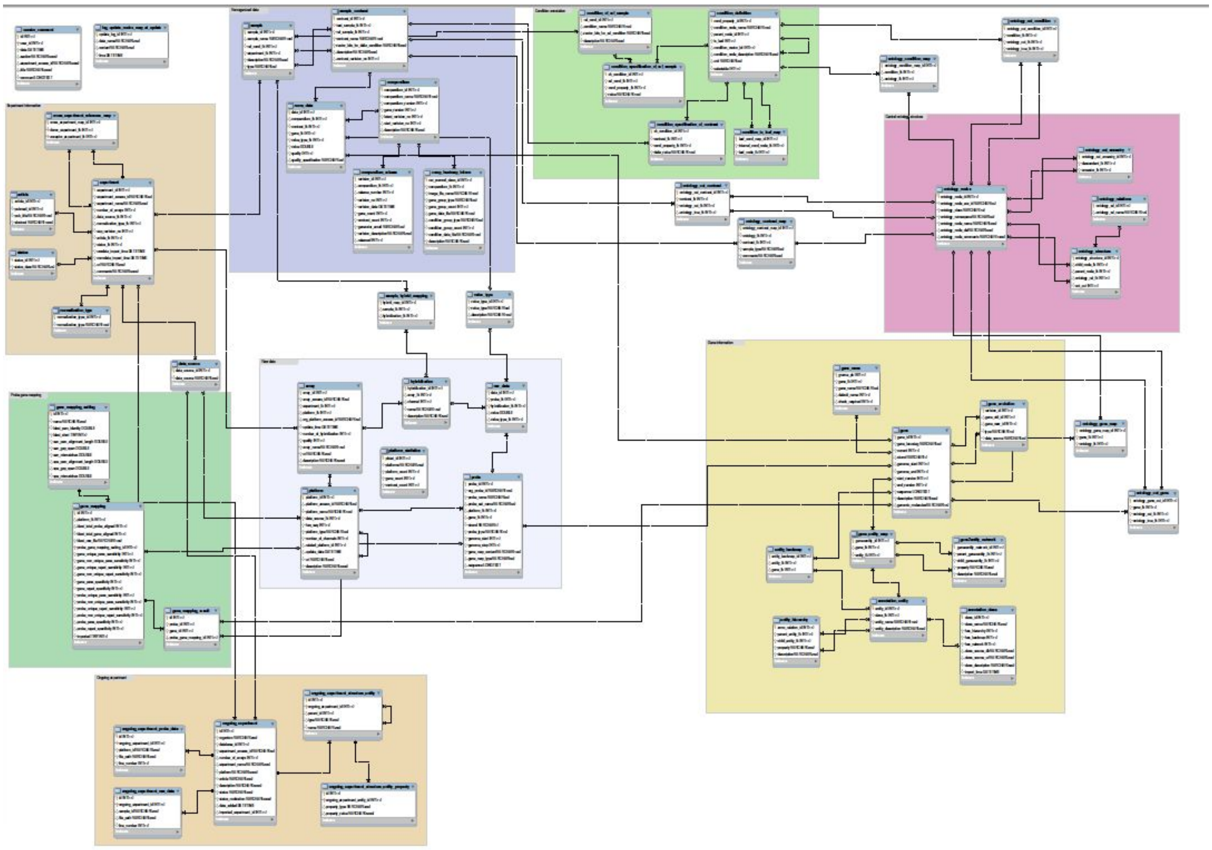

**Supplementary Figure 1.** Database schema of the MySQL created for PulmonDB. This schema contains the tables with probes, genes, samples, experiments, normalized values, raw values, etc. of PulmonDB, and each line represents the relationship with other tables; they are colored by topic. Experimental information is shown in khaki, homogenized data in purple, condition annotation in lemon green, central ontology structure in pink, probe gene mapping in green, raw data in white, gene information in yellow, ongoing experiment in wheat.

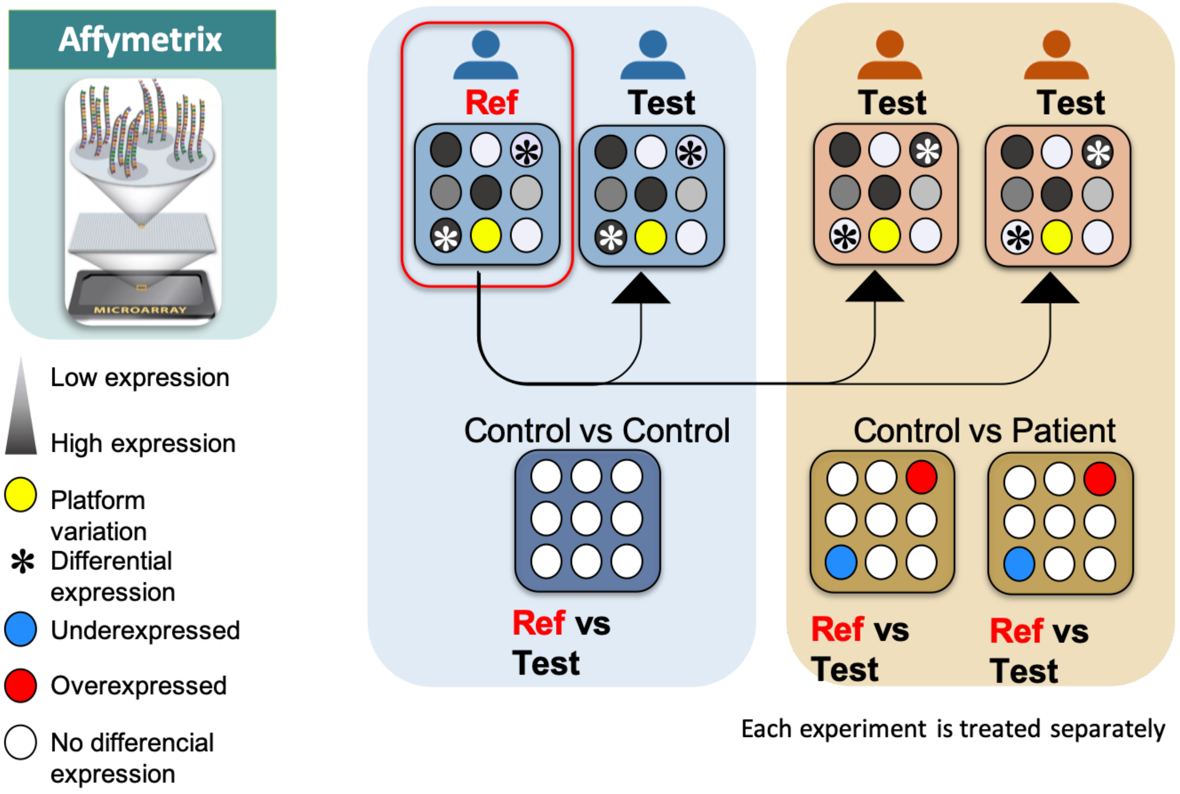


**Supplementary Figure 2**. Sample contrasts, created based on the original hypothesis of each experiment. We used a sample as a reference to make individual contrasts. The contrasts can be between control and control, control vs disease, etc. In this figure, a microarray is represented, with low expression in white and high expression in black, platform variation in yellow, differential gene expression by an asterisk, underexpressed genes in blue, overexpressed genes in red, and genes with no differential expression in white.

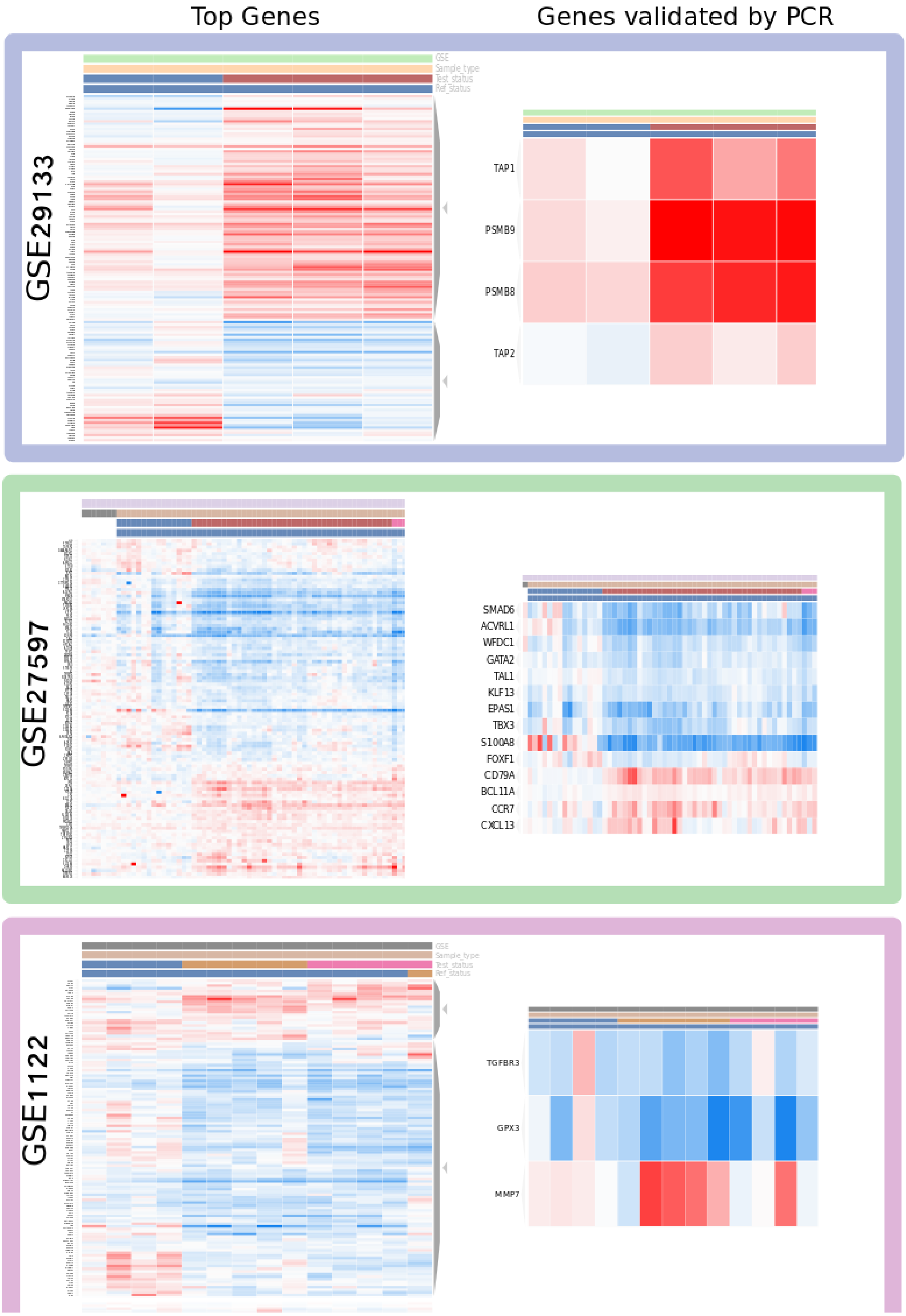


**Supplementary Figure 3.** Top differentially expressed genes in COPD published in three separate papers that also have a related GSE. Purple, genes of Fujino N, *et al.*[52] using GSE29133; green, genes of Campbell JD, *et al.*[34] using GSE27597;

pink, genes of Golpon HA, *et al.*[53] using GSE1122. The top genes and the genes validated by PCR reported in each article were used as input in PulmonDB website to check reproducibility per experiment in our homogenized contrast values.

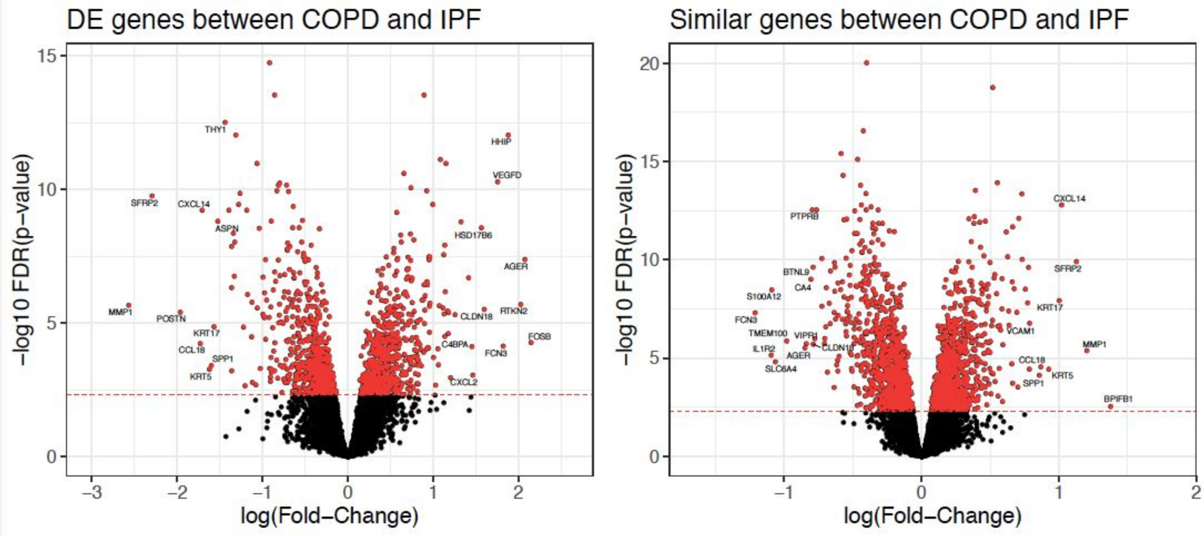


**Supplementary Figure 4.** Volcano plots of differentially expressed genes analyzed for COPD versus IPF and for COPD and IPF compared to a control group. The *x* axis is the $\log_2$ fold change, the *y* axis is the adjusted *p* -value using FDR in $-\log_{10}$, red dots are genes with an adjusted *p*-value of <0.001.

**Supplementary Table 1**. Names and descriptions of the controlled vocabulary used to annotate samples in PulmonDB.

| Name | Description | Units |
|---|---|---|
| +/-SD_AGE | Standard deviation for the group age | Boolean |
| A549 | Lung carcinoma cell line | Boolean |
| AAD | Alpha-1 antitrypsin deficiency-related emphysema | Boolean |
| ADENOVIRUS_TREATMENT | Cell culture treated with adenovirus to activate green fluorescent protein (GFP) | Boolean |
| ADVANCED | IPF advanced stage | Boolean |
| AE | Acute exacerbation | Boolean |
| AFRICAN | Ancestry of the donor: African | Boolean |
| AGE_PER_INDIVIDUAL | Age of the donor | Years |
| AIR_TREATMENT | Air treatment | Boolean |

| | | |
|---|---|---|
| AIRWAY SMOOTH MUSCLE CELLS | Airway smooth muscle cells | Boolean |
| AIRWAY-DYSPLASIA | Airway-dysplasia | Boolean |
| ALI_CULTURE | Air-liquid interface culture | Days |
| ALVEOLAR_EPITHELIAL_CELLS | Alveolar epithelial cells | Boolean |
| ALVEOLAR_MACROPHAGE | Alveolar macrophage | Boolean |
| ANTI_CD3_AND_ANTI_CD28 | Anti-CD3/CD28 cell activator | Boolean |
| ASIAN | Ancestry of the donor: Asian | Boolean |
| ASTHMATIC | Asthmatic | Boolean |
| ATRA | All-*trans* retinoic acid concentration | mM |
| AZA | 5'-Azacytidine | μM |
| BAL | Bronchoalveolar lavage (BAL) | Boolean |
| BEAS-2B | Bronchial epithelial cell line | Boolean |
| BLOOD | Blood (sample type) | Boolean |
| BONE_MARROW | Hematopoietic stem cells | Boolean |
| BRONCHIAL_EPITHELIAL_CELLS | Bronchial epithelial cells | Boolean |
| CAUCASIAN | Ancestry of donor: Caucasian | Boolean |
| CCD25Lu | Commercial fibroblast cell line | Boolean |
| CD14+ | Monocytes CD14+ | Boolean |
| CD3/28 | Anti-CD3/CD28 beads | Boolean |
| CD34+ | CD34+ hematopoietic stem cells | 0,1 |
| CD4-T-CELLS | CD4 T cells | Boolean |
| CD8-T-CELLS | CD8 T cells | Boolean |
| CIGARETTES | Number of cigarettes per year | Number_of_Cigarettes |
| CONTRACTILE_ECM | Contractile extracellular matrix | Boolean |

| | | |
|---|---|---|
| CONTROL_ECM | Control extracellular matrix | Boolean |
| COP | Cryptogenic organizing pneumonia | Boolean |
| COPD | Chronic obstructive pulmonary disease | Boolean |
| COPD_EXPERIMENT | Used to classify the disease studied in the experiment. NOTE: It does not describe whether donor is healthy or diseased. | Boolean |
| CPFE | Combined pulmonary fibrosis and emphysema | Boolean |
| CPFE_EXPERIMENT | Combined pulmonary fibrosis and emphysema; used to classify the disease studied in the experiment. | Boolean |
| CULTURED | Cultured samples | Boolean |
| CURCUMIN | Curcumin | uM |
| CURRENT | Current smoker | Boolean |
| DIAPHRAGM_MUSCLE | Diaphragm muscle | Boolean |
| DIP | Desquamative interstitial pneumonia | Boolean |
| DLCO (%) | DLCO % | % |
| EARLY | IPF early stage | Boolean |
| EBV | Epstein-Barr virus infection | |
| EFFECTOR | Differential status of T cells | 0,1 |
| EMPHYSEMA | Emphysema | Boolean |
| ENDOTHELIAL_CELLS | Endothelial cells | Boolean |
| FAMILIAL_IPF | Familial IPF sample | Boolean |
| FEMALE | Female sample | Boolean |
| FEV1/FVC (%) | FEV1/FVC (%) | % |
| FIBROBLASTS | Fibroblasts | Boolean |
| FLUTICASONE | Corticosteroid treatment for COPD | Boolean |
| FLUTICASONE_6MONTHS/PLACEBO_24MONTHS | Treatment for COPD | Boolean |
| FLUTICASONE/SALMETEROL | Corticosteroid treatment for COPD | Boolean |

| | | |
|---|---|---|
| FORMER | Former smoker, a ex-smoker individual | Boolean |
| FVC (%) | FVC percentage | % |
| GHK | Glycyl-L-histidyl-L-lysine peptide (nano molar concentration) | nM |
| GOLD_I | GOLD stage I, mild; FEV1 / FVC < 0.70;FEV1 80% predicted | Boolean |
| GOLD_II | GOLD stage II, moderate; FEV1 / FVC < 0.7050% FEV1 < 80% predicted | Boolean |
| GOLD_III | GOLD stage III, severe;FEV1 / FVC < 0.7030% FEV1 < 50% predicted | Boolean |
| GOLD_IV | GOLD stage IV, very severe;FEV1 / FVC < 0.70FEV1 < 30% predicted or FEV1 < 50% predicted plus chronic respiratory failure | Boolean |
| GROUP | Number of individuals that comprise the pool or group | individuals |
| HEALTHY/CONTROL | Normal status | Boolean |
| HFL1 | HFL1 | Boolean |
| HISPANIC | Ancestry of the donor: Hispanic | Boolean |
| HOMOZYGOUS_NULL | Homozygous null gene | null_gene |
| HOURS | Number of hours of the treatment | hrs |
| IFN_GAMMA | Interferon gamma concentration | U/mL |
| IL-13 | IL-13 | ng/mL |
| IL-6R_CD8 | IL-6R CD8 T cells | Boolean |
| IL-7R_CD8 | IL-7R CD8 T cells | Boolean |
| INDIVIDUAL | If the donor is an individual | Boolean |
| IPF | Idiopathic pulmonary fibrosis | Boolean |
| IPF_ECM | Fibrotic extracellular matrix | Boolean |
| IPF_EXPERIMENT | Used to classify the disease studied in the experiment. NOTE: It does not describe whether donor is healthy or diseased. | Boolean |
| LARGE_AIRWAY_EPITH ELIAL CELLS | Large airway epithelial cells | Boolean |

| | | |
|---|---|---|
| LATINAMERICAN | Ancestry of the donor: Latin American | Boolean |
| LEUKOCYTES | Leukocytes | Boolean |
| LIA | Leukocyte-induced angiogenesis assay | Boolean |
| LOW_BODY_MASS | Low body mass | Boolean |
| LOW_LOBE | Lower lobe | Boolean |
| LUNG_BIOPSY | Lung biopsy | Boolean |
| LUNG_CANCER | Donor with lung cancer | Boolean |
| LUNG_EXPLANT | Sample taken from a lung explant | Boolean |
| LYMPHOCYTES | Lymphocytes | Boolean |
| MACROPHAGES | Macrophages | Boolean |
| MALE | Male | Boolean |
| MALE/FEMALE_RATIO | Male/female ratio | Boolean |
| MAMMARY EPITHELIAL CELLS | Mammary epithelial cells | Boolean |
| MATCH_TISSUE_CONTROL | Control tissue with lung cancer | Boolean |
| MEAN_AGE | Average age of the group | years |
| MEMORY | Differentiation status of T cells | 0,1 |
| MESENCHYMAL_PROGENITOR_CELL | Mesenchymal progenitor cell | Boolean |
| MIDDLE_LOBE | Middle lobe | Boolean |
| MONOCYTE | Monocyte | Boolean |
| MONTHS | Number of months of the treatment | unit in months |
| MRC5 | Commercial fibroblast-like cell line MRC5 | Boolean |
| MRNA | mRNA | Boolean |
| MYELOID DENDRITIC CELLS | Myeloid dendritic cells | Boolean |
| NAIVE | Differentiation status of T cells | 0,1 |

| | | |
|---|---|---|
| NATIVE_AMERICAN | Ancestry of the donor: Native American | Boolean |
| NEUTROPHILS | Neutrophils | Boolean |
| NHLF | Fibroblast commercial cell line | Boolean |
| NON_CONTRACTILE_ECM | Non contractile extracellular matrix | Boolean |
| NON_CULTURED | Noncultured | Boolean |
| NON_IPF_ILD | Interstitial lung disease (ILD) | Boolean |
| NON_SMOKER | Nonsmoker | Boolean |
| NORMAL_BODY_MASS | Normal body mass | Boolean |
| NORMAL_RELATIVE | Non affected relative | Boolean |
| NOT_CAUCASIAN | Ancestry: Not Caucasian | Boolean |
| NSIP | Nonspecific interstitial pneumonia | Boolean |
| NTHi375 | *H. influenzae* strain NTHi375 | CFU/mL |
| OXIDATIVE_STRESS | $H_2O_2$ concentration (micromolar) | μM |
| OXYGEN_TREATMENT | Oxygen treatment | Boolean |
| P3C | Pam3Cys | μg/mL |
| PACK_PER_YEAR | No. of cigarette packs smoked per year | Number of packs per year |
| PAH | Pulmonary arterial hypertension | Boolean |
| PASSAGE (RANGE) | Range of the number of times a cell culture has been subcultured | Boolean |
| PASSAGE (SPECIFIC) | Number of times a cell culture has been subcultured | Boolean |
| PBMC | Peripheral blood mononuclear cell | Boolean |
| PGE2 | Prostaglandin E2 | μg/mL |
| PH | Pulmonary hypertension | Boolean |
| PLACEBO | Placebo | Boolean |
| POLYSOME_ASSOCIATED_RNA | Polysome associated RNA | Boolean |

| | | |
|---|---|---|
| RANGE | Group age range | Years |
| RAPID | Acute exacerbation - rapid | Boolean |
| RB-ILD | Respiratory bronchiolitis | Boolean |
| REFERENCE RNA | Stratagene Universal Human Reference RNA (catalog number 740000) | Boolean |
| SALIVA | Saliva | Boolean |
| SARCOIDOSIS | Nodular, self-limiting (N-SL) | Boolean |
| SEDENTARY | Sedentary | Boolean |
| SERUM | Serum | Boolean |
| SLOW | Acute exacerbation - slow | Boolean |
| SLPS | Standard lipopolysaccharide (LPS) | ug/mL |
| SMALL_AIRWAYS_EPITHELIAL_CELLS | Small airways epithelial cells | Boolean |
| SMALL_RNAS | Small RNAs | Boolean |
| SMOKE_EXPOSURE | Treatment with smoke exposure | Boolean |
| SPONTANEOUS_IPF | Spontaneous IPF | Boolean |
| SPUTUM | Sputum | Boolean |
| SS | Systemic sclerosis | Boolean |
| STABLE | Stable patients | Boolean |
| STATIN_USER | Statin user | Boolean |
| TGF_BETA | Exogenous TGF-beta levels | ng/mL |
| THP-1 | THP-1 cell line | Boolean |
| TIG1 | Fibroblast cell line TIG1 (Tokyo Institute of Gerontology-1) | Boolean |
| TIG7 | Embryonic pulmonary fibroblast cell line TIG7 | Boolean |
| TNF_ALFA | TNF-alpha concentration | ng/mL |
| TOTAL_RNA | Total RNA | Boolean |
| TRACHEAL EPITHELIAL CELLS | Tracheal epithelial cells | Boolean |

| | | |
|---|---|---|
| TRAINED | Trained | Boolean |
| TRAINING_EXERCISE | Training exercise (in weeks) | Number of weeks |
| TRANSFECTED | Transfection with the gene (fill in the blank with the gene name) | Gene |
| U937 | Immortalized monocyte cell line | Boolean |
| UNCLASSIFIED | Sample with unclassified disease | Boolean |
| UNTREATED | No treated cells/tissue | Boolean |
| UPLPS | Ultrapure *Escherichia coli* O111:B4 LPS | µg/mL |
| UPPER_LOBE | Upper lobe | Boolean |
| VASTUS_LATERALIS_MUSCLE | Vastus lateralis muscle | Boolean |
| WEEKS | Number of weeks of the treatment | Number of weeks |

**Supplementary Table 2**. GEO Series available in PulmonDB with the number of sample contrasts created per experiment.

| GEO Series ID | Lung disease | Experiment original name (as published) | Pubmed ID related | Platform(s) | Sample contrasts |
|---|---|---|---|---|---|
| GSE10038 | COPD | Upregulation of expression of matrix metalloproteinases in alveolar macrophages of HIV-1+ smokers with early emphysema | 19605697 | HG-U133_Plus_2 | 10 |
| GSE107426 | COPD | Results of differentially expressed lncRNAs in COPD and healthy smokers using the Arraystar Human LncRNA microarray | NA | GPL16956 | 1 |
| GSE10896 | COPD | Impact of curcumin on human monocytes (U937 cells) exposed to oxidative stress | 18421014 | HG-U133_Plus_2 | 22 |
| GSE1122 | COPD | Emphysema lung tissue gene expression profiling | 15284076 | GPL80 | 14 |
| GSE12815 | COPD | PI3K pathway activity in the normal airway of smokers with lung cancer, and in smokers with airway dysplasia | 20375364 | GPL5175,GPL96, HG-U133_Plus_2 | 67 |

| | | | | | |
|---|---|---|---|---|---|
| GSE13896 | COPD | Smoking-dependent reprogramming of alveolar macrophage polarization: implication for pathogenesis of COPD | 19635926 | HG-U133_Plus_2 | 69 |
| GSE1650 | COPD | COPD study | 15374838 | GPL96 | 29 |
| GSE16972 | COPD | COPD-specific gene expression signatures of alveolar macrophages as well as peripheral blood monocytes overlap and correlate with lung function | 21430361 | GPL96 | 11 |
| GSE2125 | COPD | Isolated alveolar macrophages | 16166618 | HG-U133_Plus_2 | 44 |
| GSE23704 | COPD | Gene expression profiling on bronchoalveolar lavage (BAL) cells treated with all-*trans*-retinoic acid (ATRA) | 22204820 | HG-U133_Plus_2 | 1 |
| GSE26296 | COPD | Genome-wide analysis of lung myeloid dendritic cells gene expression from healthy or emphysema subjects | 22261033 | GPL6884 | 5 |
| GSE27536 | COPD | Vastus lateralis biopsies from healthy and COPD patients before and after 8 weeks of exercise training | 21909251 | HG-U133_Plus_2 | 53 |
| GSE27543 | COPD | Vastus lateralis biopsies from healthy and COPD patients before and after 3 weeks of endurance training | 21909251 | GPL201 | 15 |
| GSE27597 | COPD | A gene expression signature of emphysema-related lung destruction and its reversal by the tripeptide GHK | 22937864, 24380442, 24380443, 24089408 | GPL13243 | 70 |
| GSE29133 | COPD | Transcriptome in alveolar epithelial type II cells isolated from normal and COPD lungs of adult human | 23117565 | HG-U133_Plus_2 | 5 |
| GSE30027 | COPD | Human lung CD8+ T cells compared with paired non-naive peripheral blood CD8+ T cells | NA | GPL6947 | 11 |
| GSE33337 | COPD | miRNA changes in mild and moderate emphysema correlate with target gene expression in vivo and in vitro [target gene expression data] | 24479666 | GPL6947 | 6 |

| | | | | | |
|---|---|---|---|---|---|
| GSE34562 | COPD | IL-6R identifies early differentiated human effector memory CD8+ T cells that potently expand and produce IL-2 and IL-13 | 25390970 | GPL10558 | 5 |
| GSE37147 | COPD | Bronchial airway gene expression reflects a COPD-associated field of injury that changes with disease severity and is reversible with therapy | 23471465 | GPL13243 | 269 |
| GSE37693 | COPD | Gene expression effects of IL-13 on primary human airway epithelial cells | 23187130 | GPL6947 | 7 |
| GSE37768 | COPD | Expression data in lung tissue from moderate COPD patients, healthy smokers and nonsmokers | NA | HG-U133_Plus_2 | 37 |
| GSE40885 | COPD | Data expression in alveolar macrophages induced by lipopolysaccharide in humans | 22952057 | HG-U133_Plus_2 | 13 |
| GSE45251 | COPD | The inflammatory response of human airway smooth muscle cells | 23590298 | GPL6480 | 15 |
| GSE46903 | COPD | Transcriptome-based network analysis reveals a spectrum model of human macrophage activation [Expression] | 24530056 | GPL6947 | 383 |
| GSE47460 | COPD | Gene expression profiling of chronic lung disease for the Lung Genomics Research Consortium | 29988126, 27609773, 27609774 | GPL5175, GPL13607,GPL4133 | 581 |
| GSE475 | COPD | Chronic obstructive pulmonary disease | NA | GPL96 | 6 |
| GSE47718 | COPD | Smoking dysregulates the human airway basal cell transcriptome at COPD-linked risk locus 19q13.2 | 24498427 | GPL11154 | 16 |
| GSE47929 | COPD | Gene expression profiles of Siglec-14/THP-1 and Siglec-5/THP cell lines, with or without NTHi stimulation | 24994897 | HG-U133_Plus_2 | 3 |
| GSE55962 | COPD | Systemic inflammatory response to smoking in chronic obstructive pulmonary disease: evidence of a gender effect | 24830457 | GPL13667 | 105 |
| GSE56341 | COPD | Gene expression profiles of COPD and nonCOPD small airway epithelia | 24298892 | GPL6244 | 21 |

| | | | | | |
|---|---|---|---|---|---|
| GSE56768 | COPD | Whole blood and isolated blood cell transcriptomics in COPD | NA | HG-U133_Plus_2 | 431 |
| GSE56768 | COPD | Whole blood and isolated blood cell transcriptomics in COPD | NA | HG-U133_Plus_2 | |
| GSE57148 | COPD | Characterizing gene expression in lung tissue of COPD subjects using RNA-seq | 25834810, 29871630 | GPL11154 | 186 |
| GSE60399 | COPD | Expression analysis of stable chronic obstructive pulmonary disease and acute exacerbation of chronic obstructive pulmonary disease | 25407108 | GPL18451 | 4 |
| GSE62974 | COPD | RNA sequencing (RNA-SEQ) of EPAS1 knockdown by siRNA in endothelial cells | 25569234 | GPL16791 | 5 |
| GSE63073 | COPD | Genes related to emphysema are enriched for ubiquitination pathways | 25432663 | GPL3991 | 42 |
| GSE69134 | COPD | Genome expression profiling identifies host-directed antimicrobial drugs against respiratory infection by nontypable *Haemophilus influenzae* | 26416856 | GPL887 | 7 |
| GSE69557 | COPD | $CD34^{+}$ $DNAM^{bright}$ $CXCR4^{+}$ CLP mobilized in chronic inflammation | 26436997 | GPL6244 | 3 |
| GSE69818 | COPD | COPD lung tissue expression | 26735770 | GPL13667 | 69 |
| GSE7557 | COPD | Human bronchial epithelial cells_Passage 3 vs. passage 0 | 17965775 | GPL5102 | 3 |
| GSE76705 | COPD | Complex disease subtypes identified by network-based clustering of gene expression data: application to COPD | 26773458 | HG-U133_Plus_2 | 228 |
| GSE77344 | COPD | Gene expression in chronic obstructive pulmonary disease (COPD) patients | http://dx.doi.org/10.1101/038794 | GPL11532 | 171 |
| GSE81614 | COPD | Novel RNA-binding activity of NQO1 promotes SERPINA1 mRNA translation | 27515817 | GPL10558 | 14 |
| GSE8581 | COPD | Human chronic obstructive pulmonary disorder (COPD) biomarker | 18849563 | HG-U133_Plus_2 | 58 |

| | | | | | |
|---|---|---|---|---|---|
| GSE8608 | COPD | MDM from COPD patients and healthy subjects after treatment with LPS or fine and ultrafine particles | 18084737 | HG-U133_Plus_2 | 5 |
| GSE87098 | COPD | Expression data of mucociliated human airway epithelia on-chip with or without exposure to whole cigarette smoke under physiological breathing | 27894999 | GPL16686 | 14 |
| GSE994 | COPD | Effects of cigarette smoke on the human airway epithelial cell transcriptome | 15210990 | GPL96 | 74 |
| GSE1786 | Lung function in elderly | Vastus lateralis biopsies from healthy trained and sedentary males | 16260967 | GPL96 | 22 |
| GSE101286 | IPF | Gene expression profiling of idiopathic interstitial pneumonias (IIPs): identification of potential diagnostic markers and therapeutic targets | 28821283 | GPL6947 | 14 |
| GSE11196 | IPF | Fibrotic myofibroblasts manifest genome-wide derangements of translational control | 18795102 | HG-U133_Plus_2 | 44 |
| GSE15197 | IPF | RNA expression profiling of lung tissue identifies mutually distinct molecular signatures in PAH and PH secondary to IPF | 20081107 | GPL6480 | 38 |
| GSE19976 | IPF | Gene expression analysis of lung biopsies from patients with two different forms of pulmonary sarcoidosis | 20194811 | GPL6244 | 14 |
| GSE21369 | IPF | Gene expression profiles of interstitial lung disease (ILD) patients | 21241464 | HG-U133_Plus_2 | 28 |
| GSE24206 | IPF | Validated gene expression signatures of idiopathic pulmonary fibrosis | 21974901 | HG-U133_Plus_2 | 22 |
| GSE26594 | IPF | Increased cell surface Fas expression is necessary to sensitize lung fibroblasts to Fas ligation-induced apoptosis: implications for fibroblast accumulation in idiopathic pulmonary fibrosis | 21632719 | HG-U133_Plus_2 | 5 |

| | | | | | |
|---|---|---|---|---|---|
| GSE28221 | IPF | Peripheral blood mononuclear cell gene expression profiles may predict poor outcome in idiopathic pulmonary fibrosis | 24089408 | GPL4133,GPL5175 | 136 |
| GSE31934 | IPF | To examine the expression of Sulf1 and Sulf2, as well as other glycan-related genes, in human Idiopathic pulmonary fibrosis (IPF) lungs compared to normal lung samples | NA | GPL11097 | 5 |
| GSE32537 | IPF | Molecular phenotyping of the idiopathic interstitial pneumonias [mRNA] | 23783374 | GPL6244 | 216 |
| GSE33566 | IPF | The peripheral blood transcriptome predicts the presence and extent of disease in idiopathic pulmonary fibrosis | 22761659 | GPL4133 | 122 |
| GSE35145 | IPF | Gene expression changes in IPF | 22700861 | GPL10558 | 7 |
| GSE38958 | IPF | Profiling of gene expression in idiopathic pulmonary fibrosis | 26286721 | GPL5175 | 114 |
| GSE44723 | IPF | Bleomycin induces molecular changes directly relevant to idiopathic pulmonary fibrosis: A model for “active” disease | 23565148 | HG-U133_Plus_2 | 12 |
| GSE45686 | IPF | An extracellular matrix-driven positive feedback loop regulates translation in idiopathic pulmonary fibrosis | 24590289 | GPL10558 | 38 |
| GSE48149 | IPF | Lung tissues in systemic sclerosis have gene expression patterns unique to pulmonary fibrosis and pulmonary hypertension | 21360508 | GPL16221 | 52 |
| GSE49072 | IPF | Alveolar macrophage gene expression in human pulmonary fibrosis | 23924348 | GPL96 | 83 |
| GSE52463 | IPF | Transcriptome analysis reveals differential splicing events in IPF lung tissue | 24647608 | GPL11154 | 14 |
| GSE52612 | IPF | Forkhead Box F1 (FOXF1) represses fibroblast functions relevant to fibrogenesis | 25260753 | GPL13607 | 7 |
| GSE53845 | IPF | 40 IPF patients and 8 healthy controls | 25217476 | GPL4133 | 94 |

| | | | | | |
|---|---|---|---|---|---|
| GSE5457 | IPF | Retinoic acids exposure alters TGF-beta1-induced epithelial mesenchymal transition via Wnt5b expression | 18621908 | GPL96 | 7 |
| GSE6804 | IPF | Expression data of myofibroblast isolated from patients with idiopathic pulmonary fibrosis | 17986007 | GPL201 | 2 |
| GSE69764 | IPF | Identification of the gene expression in IPF lung fibroblasts after demethylation | 26442443 | HG-U133_Plus_2 | 5 |
| GSE71351 | IPF | Global gene expression profiles of fibroblasts from the lungs of patients with idiopathic pulmonary fibrosis: The role of CCL8 | 28057004 | GPL10558 | 11 |
| GSE72073 | IPF | Expression data from lung tissues of IPF patients and normal controls | 26453058 | GPL17586 | 7 |
| GSE73854 | IPF | Developmental programming in idiopathic pulmonary fibrosis (IPF) | 27869174 | HG-U133_Plus_2 | 7 |
| GSE94060 | IPF | Human lung MPC | 28463231 | GPL6244 | 8 |
| GSE38934 | IPF/COPD | Gene expression profiling of lung tissues from patients with combined pulmonary fibrosis and emphysema | 23025845 | HG-U133_Plus_2 | 5 |
